# Inter-slice leakage and intra-slice aliasing in simultaneous multi-slice echo-planar images

**DOI:** 10.1101/458042

**Authors:** Carolyn Beth McNabb, Michael Lindner, Shan Shen, Laura Grace Burgess, Kou Murayama, Tom Johnstone

**Author notes:** CM and ML contributed equally to this work. Corresponding author: Dr Carolyn Beth McNabb, School of Psychology and Clinical Language Sciences, University of Reading, Harry Pitt Building, Earley Gate, Reading RG6 7BE, United Kingdom. (ext 7937). Contributions: All authors contributed to the study conception and design. Material preparation, data collection and analysis were performed by Carolyn McNabb and Michael Lindner. Shan Shen assisted with SMS sequence selection and scanning. The first draft of the manuscript was written by Carolyn McNabb. All authors commented on previous versions of the manuscript and have read and approved the final manuscript.

## Abstract

Simultaneous multi-slice (SMS) imaging is a popular technique for increasing acquisition speed in echo-planar imaging (EPI) fMRI. However, SMS data are prone to motion sensitivity and slice leakage artefacts, which spread signal between simultaneously acquired slices. Relevant to motion sensitivity, artefacts from moving anatomic structures propagate along the phase-encoding (PE) direction. This is particularly relevant for eye movement. As signal from the eye is acquired along with signal from simultaneously excited slices during SMS, there is potential for signal to spread in-plane and between spatially remote slices. After identifying an artefact temporally coinciding with signal fluctuations in the eye and spatially distributed in correspondence with multiband slice acceleration and parallel imaging factors, we conducted a series of small experiments to investigate eye movement artefacts in SMS data and the contribution of PE direction to the invasiveness of these artefacts. Five healthy adult volunteers were scanned during a blinking task using a standard SMS-EPI protocol with posterior-to-anterior (P≫A), anterior-to-posterior (A≫P) or right-to-left (R≫L) PE direction. The intensity of signal fluctuations (artefact severity) was measured at expected artefact positions and control positions. We demonstrated a direct relationship between eye movements and artefact severity across expected artefact regions. Within-brain artefacts were apparent in P≫A- and A≫P-acquired data but not in R≫L data due to the shift in artefact positions. Further research into eye motion artefacts in SMS data is warranted but researchers should exercise caution with SMS protocols. We recommend rigorous piloting of SMS protocols and switching to R≫L/L≫R PE where feasible.

## 1. Introduction

Simultaneous multi-slice (SMS (Larkman et al. 2001; Setsompop et al. 2012), multi-band; MB) techniques for functional magnetic resonance imaging (fMRI) substantially reduce the acquisition time of echo-planar imaging (EPI) data (Nunes et al. 2006), increase temporal and spatial resolution and improve statistical results of functional network analyses (Preibisch et al. 2015; Demetriou et al. 2018).

Despite rigorous piloting of SMS parameters for large-scale projects (e.g. the Human Connectome Project; HCP), optimal parameters for smaller-scale, often time-limited studies have not been established. When scanning time is limited, HCP recommends using A≫P or P≫A phase-encoding for single resting-state or task-fMRI runs to avoid right/left susceptibility asymmetry (bias) in the aggregate data caused by R≫L or L≫R phase-encoding (PE) (Human Connectome Project 2013).

However, adjusting parameters such as the PE direction can be detrimental to SMS image quality. PE direction influences the direction of susceptibility, flow, motion artefacts, and also determines the direction of aliasing that can lead to “apparent” activation distilled from actual activated brain regions. Artefacts from moving anatomic structures and signal dropout from air-tissue interfaces all propagate along the PE direction. One such anomaly is caused by eye movement. Chen and Zhu (1997) recommend acquiring data along the P≫A direction or employing saturation bands around the eyes to avoid artefacts caused by eye movement. With SMS, however, as signal from the eye is acquired along with signal from simultaneously excited slices, there is potential for signal to spread not only in-plane but between spatially remote slices (Todd et al. 2016). This is referred to as slice-leakage. Although recent methods, such as split slice-GRAPPA (Cauley et al. 2014), have been developed to reduce slice-leakage artefacts, under certain conditions, they still occur.

In this short communication, we introduce an artefact that is unique to SMS-EPI data, caused in this case by eye movement during image acquisition, but that has broader potential implications concerning more subtle artefacts resulting from real BOLD activation. The artefact was first identified while acquiring P≫A-encoded SMS data for our two recent studies; it covered parts of the temporal lobes, frontal pole/cerebral white matter, lateral occipital cortex and precentral/superior frontal gyri in some individual subjects’ data. The temporal signature of the artefact corresponded with signal fluctuations in the eye and the in-plane component resembled eye motion artefacts previously described by others (Chen and Zhu 1997). The spatial configuration of the artefact was consistent between individuals and coincided with the predicted arrangement based on SMS and GRAPPA (GeneRalized Autocalibrating Partial Parallel Acquisition) parameters used to acquire the data (Todd et al. 2016). Hypothesising that the issue was caused by signal aliasing from the eye, we were able to replicate the artefact in a healthy adult subject by asking them to blink forcefully. This procedure produced periods of signal fluctuation in expected artefact regions that discontinued during eyes-open rest (Fig. 1 and Online Resource S1). Although in-plane aliasing from eye movement is a well-known phenomenon in neuroimaging, the substantial added effect of multiband acceleration on this aliasing has not been addressed. We conducted a series of small experiments to investigate eye movement artefacts in SMS data and the contribution of PE direction to the invasiveness of these artefacts.

**Fig. 1.**
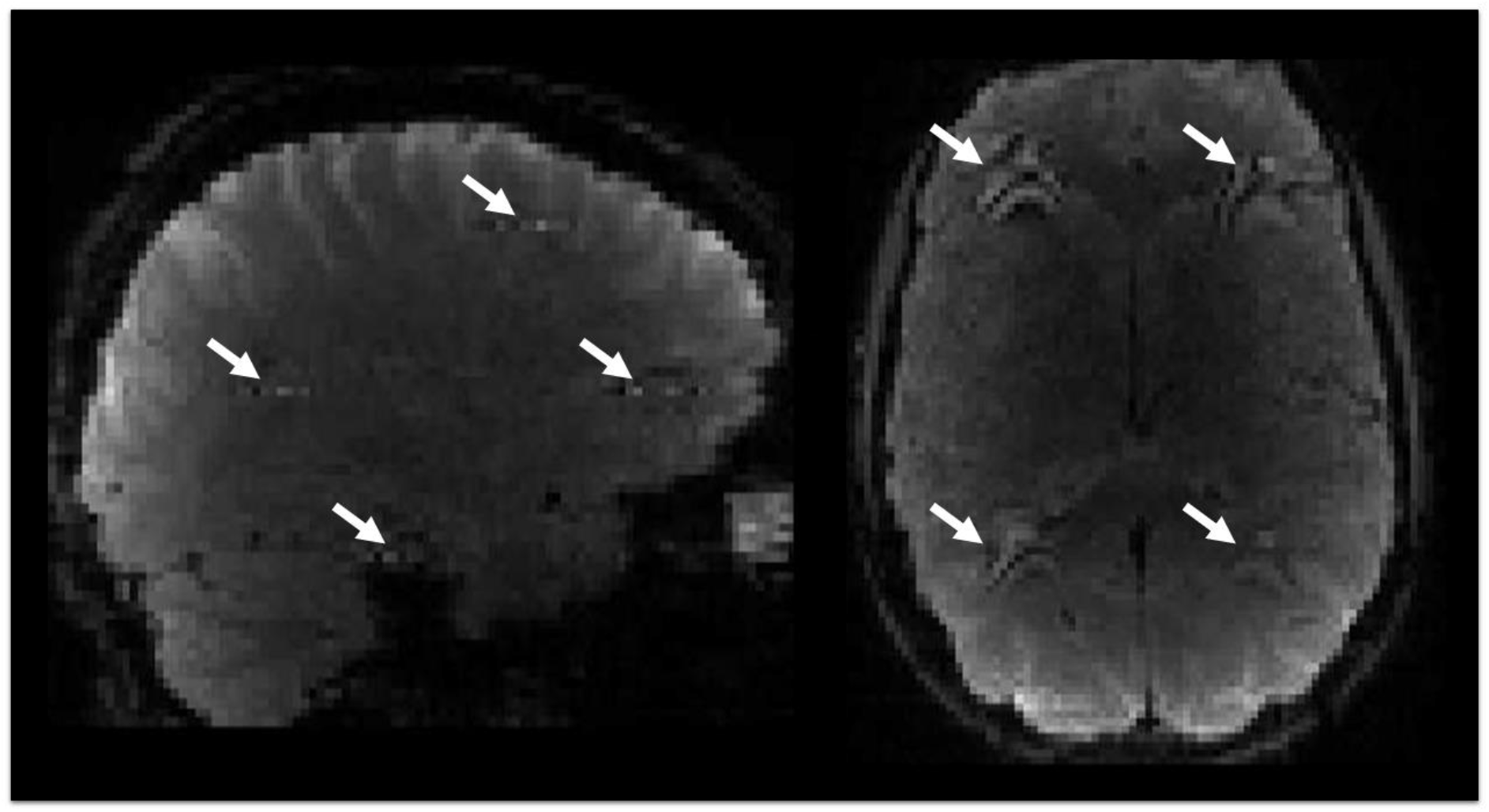
Single volume displaying artefact in individual subject. Dynamic (4 dimensional) time series data are provided in Online Resource S1

## 2. Method

Five healthy adult volunteers (3 male, 2 female) were scanned during a simple blinking task, consisting of 7.5 alternating blocks of 20 s forceful blinking and 20 s rest (eyes open blinking naturally), total scan time 5 minutes. The protocol was approved by the University of Reading Research Ethics Committee. All participants provided written informed consent.

Data were acquired on a Siemens MAGNETOM Prisma^fit^ 3T MRI scanner (Siemens, Erlangen, Germany) using a 32-channel radiofrequency head coil and the Siemens SMS BOLD (two-dimensional (2D) multiband gradient echo EPI (SMS-EPI)) sequence, optimized for the 32-channel coil. Acquisition parameters were as follows: 2 × 2 mm voxels in-plane; 2 mm slice thickness with 0% slice gap; 68 slices; 192 × 192 mm in-plane field-of-view (FOV); repetition time (TR) = 1.5 s; echo time (TE) = 30 ms; effective echo spacing 0.47 ms; GRAPPA 2 in-plane; fat saturation, PE direction P≫A, MB slice acceleration (MB4).

### Study A

To demonstrate that the artefact is reproducible, MB4 SMS data were collected from 4 participants using the P≫A PE direction. To illustrate the robustness of SMS-related artefacts from eyelid movements, we evaluated SMS data from these 4 subjects collected using additional combinations of phase encode (PE) direction and multiband slice acceleration factor. Combinations included PE direction A≫P with multiband slice acceleration factor 4 (reverse PE of that used in the main analysis) and PE direction P≫A with multiband slice acceleration factor 2 (same PE with lower multiband factor compared with that used in main analysis). The Siemens SMS sequence employs a slice-GRAPPA reconstruction technique to tease apart signal from simultaneously acquired slices. Given that others have shown that split slice-GRAPPA can reduce displacement of activation in SMS data compared with slice-GRAPPA (Todd et al. 2016), we also sought to determine whether use of split slice-GRAPPA k-space reconstruction could prevent artefacts from forceful eye blinks. For this, we employed the multiband EPI sequence provided by the Center for Magnetic Resonance Research (CMRR) (Moeller et al. 2010). Details are provided in the online resources.

### Study B

As the location of the artefact is dependent on the FOV and in-plane angle (Todd et al. 2016), we collected SMS data from one subject in the R≫L PE direction using both standard and tilted FOV, anticipating that this would reduce the impact of eyelid movement in the brain and instead shift the artefact laterally. Two scans were acquired in this subject using R≫L PE direction, one with a standard FOV and one with FOV tilted to match HCP data acquisition parameters (T > C-20.0 (Human Connectome Project 2013)), a third scan was acquired using P≫A PE direction. All other parameters remained the same as previous scans.

Signal variance over time (i.e. the total timeseries) was calculated for each voxel in the image (pyfMRIqc available from: https://github.com/DrMichaelLindner/pyfMRIqc); regions associated with the artefact were expected to show greater variance compared with unaffected regions. Artefact locations were detected using in-house-designed Matlab software (Mathworks, Natick, Massachusetts, USA), MAP4SL (available from: https://github.com/DrMichaelLindner/MAP4SL). Expected artefact locations (see Fig. 2a) were determined based on Controlled Aliasing in Parallel Imaging (CAIPI)-related FOV and in-plane GRAPPA shifts associated with the SMS sequence (Todd et al. 2016). Within each simultaneously acquired slice, two alias locations were expected: one for the CAIPI shift ((FOV/3)*m) and one for GRAPPA ((FOV/3)*m + FOV/2, where M is the number of simultaneously acquired slices and m goes from 1 to M) (Todd et al. 2016). Two control regions were located within the brain spatially isolated from expected artefact regions. A 29-voxel, in-plane mask with a diameter of 7 voxels was created for each artefact location.

**Fig. 2.**
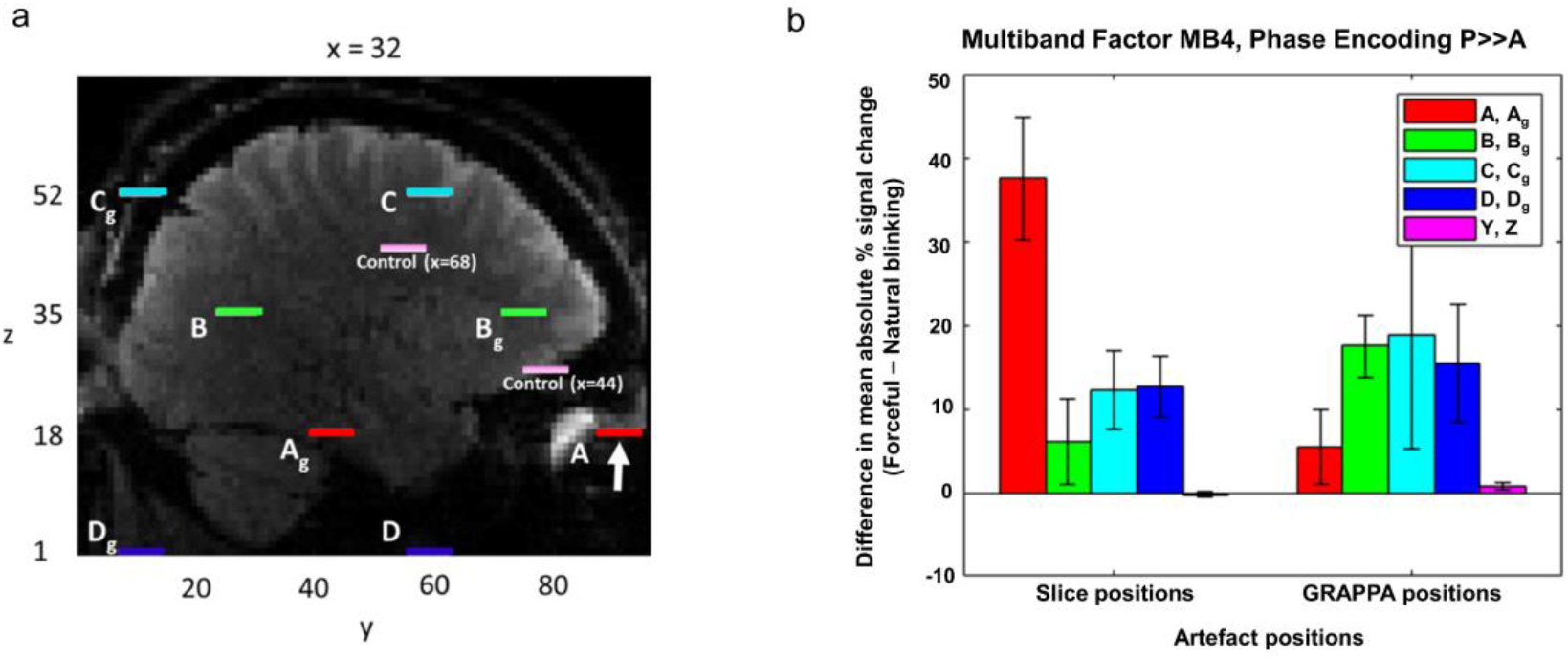
a) Expected artefact locations in right hemisphere for an individual subject (centres of expected artefact disks are shown). A-D represent artefact positions expected based on SMS slice acceleration factor (MB4) and CAIPI shift (FOV/3); A_g_-D_g_ represent artefact positions expected based on parallel imaging factor (GRAPPA-2; ((FOV/3)*m + FOV/2), where m is the number of simultaneously acquired slices. Coordinates for artefact source: x=32, y=91, z=18; indicated by white arrow. Control regions are shown in light pink: y and z coordinates are as shown in the figure; x coordinates are specified for each control mask. b) Difference (±standard error of the mean; SEM, n=4 subjects) in signal fluctuation between forceful blinking (on) and natural blinking (off) blocks; data are shown for expected artefact and control regions (right hemisphere). Control regions X and Y correspond to mean ± standard deviation [x y z] coordinates (in subject space) of [43±1.9 65±9.6 27±4.0] and [67±1.9 41±9.6 44±4.0], respectively.

The artefact is event-locked to eyelid movement, whereby shutting the eyelids elicits signal distortions and opening the eyelids results in signal recovery. Periods of forceful blinking result in large repeated changes in signal intensity at artefact positions. Volume-to-volume signal fluctuations were used to evaluate the intensity of the artefact during forced blinking and natural blinking blocks (Chen and Zhu 1997). For each masked voxel, the volume-to-volume signal change (i.e. the change in signal at each voxel between two consecutive volumes) was divided by the maximum volume-to-volume signal change (i.e. the maximum volume-to-volume change for that voxel across the entire timeseries), yielding a percentage signal change. Mean absolute percentage signal change across all voxels in the mask (29-voxel disk) was determined separately for forceful blinking and natural blinking blocks. The intensity of the artefact at each mask location was then quantified as the difference in mean absolute signal change between forceful blinking and natural blinking blocks.

## 3. Results

### Study A

Areas of highest signal variance in the brain coincided with expected artefact locations in both hemispheres, as determined by the slice acceleration factor, CAIPI shift and parallel imaging (GRAPPA) factor (Fig. 2a); whole brain signal variance for an individual subject is depicted in the left column of Fig. 3. All subjects demonstrated similar patterns of artefact intensity across forceful blinking and natural blinking blocks. Mean absolute signal change was greatest during forceful blinking blocks compared with natural blinking blocks for all expected artefact regions but not for control regions (Fig. 2b). The artefact was not limited to the sequences measured above and was also present (though to a lesser extent) at expected locations in data collected using MB slice acceleration factor MB2 (intraslice artefact only) and PE direction A≫P (Online Resource, Fig. S2). This was expected based on projected artefact locations in both instances. Standard motion correction had a limited effect on artefact intensity (i.e. volume-to-volume signal fluctuations were reduced but the artefact was still visually apparent, see Fig. S3 and Fig. S4), as did independent components analysis (ICA)-based motion artefact correction (Fig. S4).

**Fig. 3.**
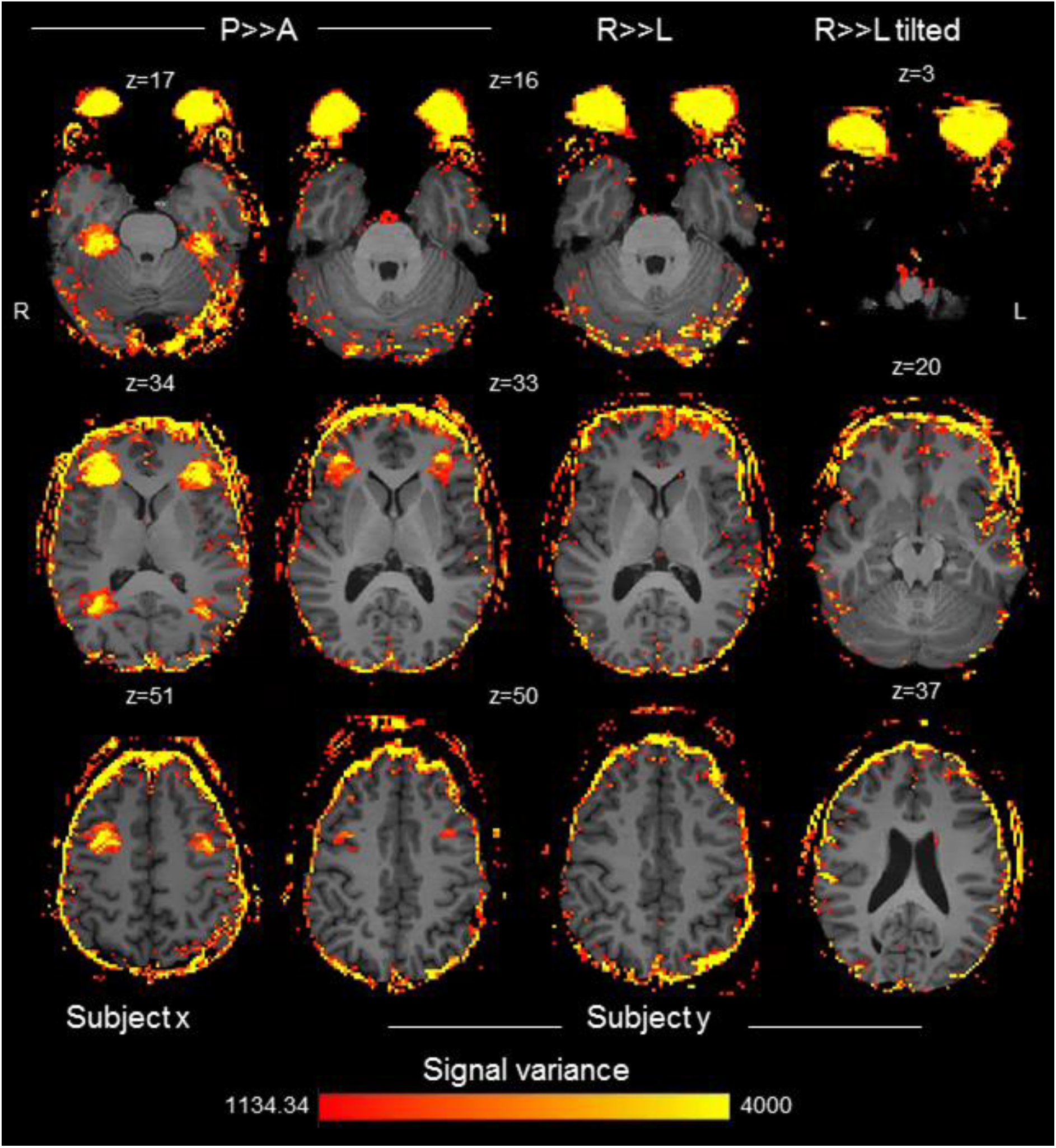
Signal variance within expected artefact slices acquired using PE direction P≫A (left columns), R≫L (middle right column) and R≫L tilted FOV (far right column). P≫A-encoded data are presented for two subjects to illustrate the replicability of the artefact

### Study B

Data for P≫A, R≫L and R≫L tilted acquisitions are shown in Fig. 3. Adjusting the PE direction to R≫L reduced in-brain artefacts in both tilted and standard FOV positions compared with P≫A data. Only a small region of the frontal pole exhibited some artefact distortion. Distortion occurred in expected artefact locations determined by the CAIPI shift and parallel imaging factor, the majority of which were located outside of the brain for R≫L acquisitions.

## 4. Discussion

SMS-EPI provides an unparalleled advantage over traditional EPI techniques in terms of its temporal resolution; however, users should exercise caution when adjusting the PE direction from the well-established R≫L or L≫R. Data presented here demonstrate that in SMS data acquired in the P≫A and A≫P PE directions, artefacts caused by eye movement leak into simultaneously acquired slices at positions predetermined by the multiband slice acceleration factor and in-plane acceleration factor. Although artefacts were less severe in data acquired using PE direction A≫P, the presence of any artefact within the brain is of major concern and should be avoided if possible. In addition, acquisition of fMRI data using the A≫P rather than P≫A PE direction compresses (rather than stretches) signal from frontal regions (e.g. orbitofrontal cortex) which is then more difficult to recover. Therefore, P≫A PE direction may be preferable for studies focussing on this region (Mori et al. 2018). Findings from this research are important for many ongoing studies using P≫A/A≫P phase-encoding with SMS-EPI, whereby researchers are unaware of the potentially detrimental impacts of SMS on image quality.

Slice leakage arises from inadequate separation of simultaneously acquired slices during k-space reconstruction. This is possibly due to poor correspondence between training (auto-calibration scans) and real-time data (Setsompop et al. 2012), as occurs in the case of subject motion (Kelly et al. 2013). In the current study, lower signal intensity fluctuations in control regions suggest that head motion alone is not responsible for the artefact reported herein. In addition, standard and ICA-based motion correction techniques (see Online Resources) were unable to eliminate the artefact in the current study, whereas motion effects reported in previous literature were successfully removed from SMS data using ICA-based approaches (Kelly et al. 2013). Standard fMRI motion correction works through whole-brain registration of each volume with a reference volume, and is designed to correct for overall movement of the head. It is not effective for correcting motion in small parts of the head, such as the eyes. In contrast, ICA is designed to remove artefacts that correspond to discrete events, such as eyeblinks, or events when such artefacts occur only in discrete regions. However, the ineffectiveness of ICA correction in this instance suggests that SMS eyeblink artefacts exhibit a different form of signal disruption. Upon visual inspection, within-brain artefacts caused by forceful blinking contained both grey and black voxels; it is therefore possible that these artefacts contain a mixture of true signal from the brain and signal drop-out from the eye region. As motion correction techniques are unable to recover lost signal, this could explain how both standard and ICA-based motion correction techniques failed to adequately correct signal in these areas.

A full-scale systematic investigation evaluating how each parameter affects the severity of non-neuronal artefact leakage is required. Based on the limited number of experiments included in this study, we cannot make any firm recommendations regarding acquisition of SMS-EPI data. However, adjustment of the PE direction to acquire data R≫L was able to minimise the impact of leakage from eye motion even at the single subject level, suggesting that this may be a more desirable PE direction for those embarking on future SMS studies. Although the artefact was minimised using R≫L phase encoding, right/left susceptibility asymmetry, resulting in compression of right frontal areas, was identified. Susceptibility distortions were left uncorrected in the current study to demonstrate that the artefact was not due to any processing stage after k-space reconstruction; however, susceptibility bias may be corrected using paired acquisition in the L≫R PE direction and subsequent distortion correction (Holland et al. 2010), as implemented in the Human Connectome Project (Human Connectome Project 2013), though this might require lengthened scan protocols. For those wishing to acquire a single fMRI run or multiple runs with the same PE direction, however, HCP recommends using P≫A or A≫P encoding (Human Connectome Project 2013). In such cases, adjustment of the multiband slice acceleration factor (to ensure that areas of interest are not collected simultaneously with regions of high signal variation - our MB2 data demonstrated only in-plane artefacts) or removal of in-plane acceleration (GRAPPA) from the protocol may reduce the risk of artefacts from eye motion. We recommend thorough piloting of all SMS-EPI sequences prior to data collection and offer researchers a tool to identify potential artefact positions (MAP4SL available from: https://github.com/DrMichaelLindner/MAP4SL).

Artefacts from slice leakage are overtly apparent in this case due to the large signal fluctuations associated with eye movement; however, more insidious inter- and intra-slice aliasing will result from any signal in the image, resulting in apparent activations reproduced and displaced from actual activated regions. Use of split slice-GRAPPA may be effective for reducing rates of false positive activation but does not adequately mitigate leakage of more severe signal changes (see Fig. S5). These issues are sufficient to warrant further investigation and should be considered before embarking on research employing SMS-EPI.

## Acknowledgments

The authors would like to thank the volunteers who gave their time to take part in the study. We also wish to that the Centre for Integrative Neuroimaging and Neurodynamics for their support throughout the project.

The study was partly supported by JSPS KAKENHI (15H05401, 16H06406, and 18H01102 to K.M.), F. J. McGuigan Early Career Investigator Prize (to K.M.), and the Leverhulme Trust (RPG-2016-146 and RL-2016-030 to K.M.).

