## Supplementary for "Inter-slice leakage and intra-slice aliasing in simultaneous multi-slice echo-planar images"

Online Resources

1. Supplementary methods

To illustrate the robustness of simultaneous multi-slice (SMS)-related artefacts from eyelid movements, we evaluated SMS data from 4 subjects collected using either phase encoding (PE) direction A>>P or multiband slice acceleration factor 2. Artefact locations were determined individually for each MR sequence and each subject. When no artefact was visible, the expected artefact location was selected based on other data for that subject. For MB2 data, the seed was placed in the eye region, as no artefact was expected in the area used for other SMS sequences (i.e. near the frontal orbital and insular cortices).

We evaluated a multiband slice acceleration factor of 2 to test the hypothesis that artefact locations are influenced by multiband slice acceleration and GRAPPA factors (as described by (Todd et al., 2016)). Decreasing the MB slice acceleration factor from MB4 to MB2 should reduce the number of slices affected by the artefact. Therefore C and $C_g$ artefacts for MB2 will be in the slice associated with C and $C_g$ in the MB4 data; however, no artefacts would be expected in the slice associated with B and $B_g$ in the MB4 data. For MB2 data, we used the standard Siemens SMS echo-planar imaging (EPI) sequence, optimized for the 32-channel coil. Parameters were as follows: 2 × 2 mm voxels in-plane; 2 mm slice thickness with 0% slice gap; 68 slices; 192 × 192 mm in-plane field-of-view (FOV); echo time (TE) = 30 ms; GRAPPA 2 in-plane; PE direction P>>A; fat saturation and MB acceleration factor 2 (MB2, repetition time (TR) = 2.25 s, echo spacing 0.38 ms).

Artefacts from eye movement spread along the PE direction (Chen and Zhu, 1997). Switching the PE direction from P>>A to A>>P will therefore influence the direction of signal leakage. However, as the direction of signal leakage still lies within the same plane, the artefact will likely still be present. We evaluated artefact severity in the Siemens MB4 sequence (described in main text) using PE direction A>>P.

To exclude the possibility that the artefact was caused by head motion rather than eye blinks, we performed standard motion correction on Siemens MB4 data and compared the difference in absolute signal change between forceful blinking and natural blinking periods. Processing of fMRI data was carried out using FEAT (FMRI Expert Analysis Tool) Version 6.00, part of FSL (FMRIB's Software Library, www.fmrib.ox.ac.uk/fsl). The following pre-statistics processing was applied; motion correction using MCFLIRT (Jenkinson et al., 2002); grand-mean intensity normalisation of the entire 4-dimensional dataset by a single multiplicative factor; and highpass temporal filtering (Gaussian-weighted least-squares straight line fitting, with sigma=50.0s).

Retrospective “clean-up” of SMS-EPI data with independent components analysis (ICA)-based motion correction has been shown to successfully remove artefacts in previous work (Kelly et al., 2013). We examined whether ICA-based motion correction could successfully remove the artefact we observe when participants are asked to blink heavily. ICA-based artefact detection was carried out using Multivariate Exploratory Linear Optimized Decomposition into Independent Components (MELODIC), part of FSL (Beckmann and Smith, 2004; Jenkinson et al., 2012). In order to preserve the structure of the artefact, no spatial smoothing was applied to the data. Data de-noising was performed by regressing out artefact components hand-selected for visual (spatial) resemblance to the artefact.

Previous work by (Todd et al., 2016) and (Cauley et al., 2014) suggests that a split slice-GRAPPA approach to k-space reconstruction can reduce unaliasing errors in SMS data compared with slice-GRAPPA reconstruction and reduce displacement of detected activation. We therefore sought to evaluate whether split slice-GRAPPA reconstruction can minimise SMS artefacts caused by forceful eye blinks. We opted to test this method using the Minnesota CMRR (Center for Magnetic Resonance Research) multiband sequence (Moeller et al., 2010; Setsompop et al., 2012; Xu et al., 2013), as split slice-GRAPPA (or leak block) can easily be turned ‘on’ or ‘off’ at the discretion of the MR technician. All parameters for the CMRR sequence were the same as those mentioned above for the Siemens MB4 sequence but with echo spacing 0.32 ms. To investigate the implications of leak block, we acquired two sets of data using the CMRR sequence, one using slice-GRAPPA reconstruction (leak block off) and one using split slice-GRAPPA reconstruction (leak block on).

### 1. Supplementary results

A video (MP4 format) of the raw SMS-EPI data illustrating slices with expected artefacts is provided in S1. Results of the MB2-acquired data are shown in Fig. S2a. Decreasing the multiband acceleration factor to MB2 reduced interslice artefact intensity, though intraslice artefacts were still present ($A_g$ region). In contrast, interslice GRAPPA-related artefacts were still present after adjusting the PE direction of MB4 data to A>>P (see Fig. S2b).

Standard motion correction techniques reduced artefact intensity but did not diminish it entirely (Fig. S3), suggesting that the artefact is not dependent on head motion and is in fact related to eye/eyelid movement. Minimal improvement in artefact severity was observed with ICA-based noise correction; results are shown in Fig. S4.

Split slice-GRAPPA reconstruction produced smaller artefacts than slice-GRAPPA reconstruction using the Minnesota CMRR multiband sequence (see Fig. S5). Though not as severe as those observed with slice-GRAPPA reconstruction, intra- and interslice artefacts seen with split slice-GRAPPA reconstruction may still interfere with signals of interest throughout the brain.

---

**Video S1** Video of artefact-affected slices from an individual subject

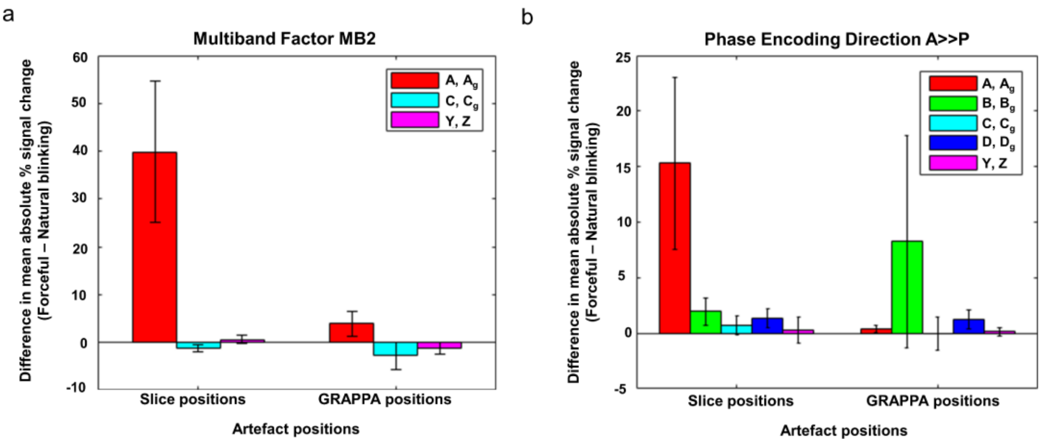


**Fig. S2** Mean difference (±SEM, n=4 subjects) in mean absolute signal change between blinking on and off blocks for expected artefact and control regions (right hemisphere) for a) multiband slice acceleration factor MB2 with PE direction P>>A (control regions X and Y correspond to mean ± standard deviation (SD) [x y z] coordinates (in subject space) of [56±1.1 45±3.4 34±3.3] and [80±1.1 17±9.1 51±3.3], respectively) and b) multiband slice acceleration factor MB4 with PE direction A>>P (control regions X and Y correspond to mean ± SD [x y z] coordinates of [44±0.8 67±8.1 28±2.9] and [68±0.8 43±8.1 45±2.9], respectively)

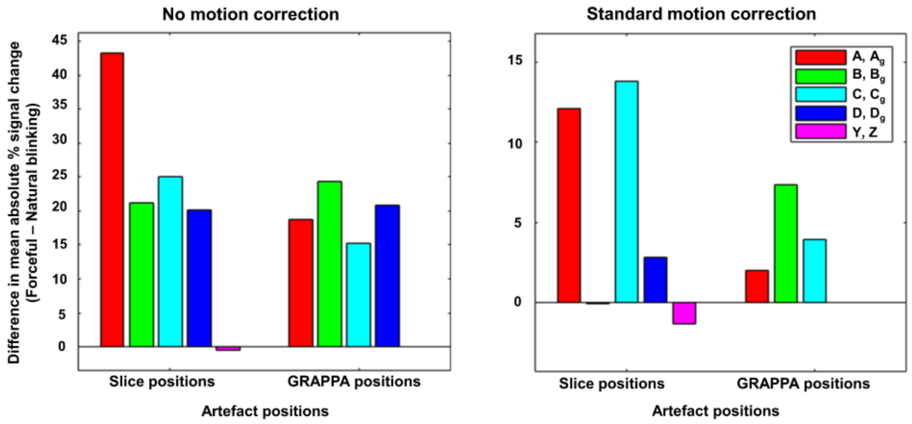


**Fig S3** Difference in mean absolute signal change between blinking on and off blocks for expected artefact and control regions across individual subject (right hemisphere) before (left) and after (right)

standard motion correction and intensity normalisation (Siemens SMS sequence). Control regions X and Y correspond to [x y z] coordinates (in subject space) of [45 54 43] and [69 30 60] respectively

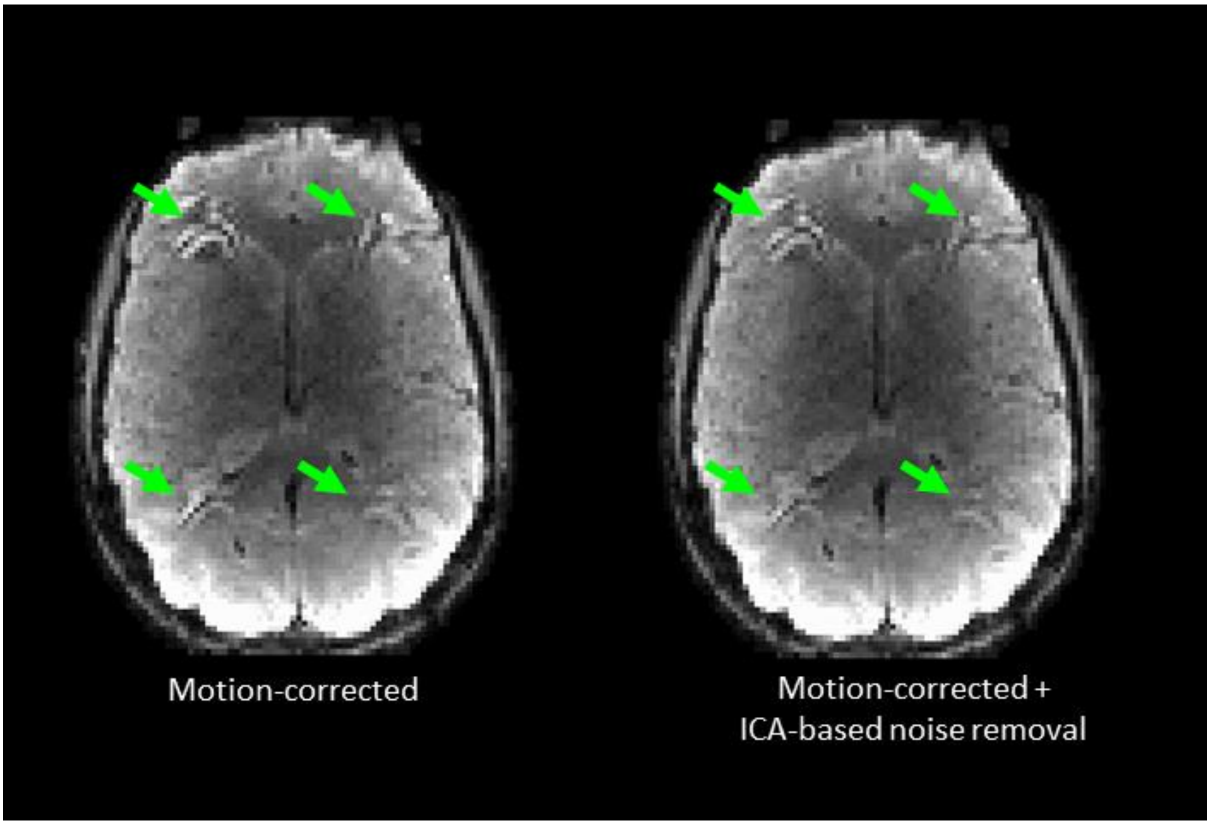


**Fig. S4** Example volume of single subject BOLD data during blinking block following standard motion correction (left) and motion correction plus ICA-based noise removal (right; Siemens SMS sequence). Arrows indicate artefact locations

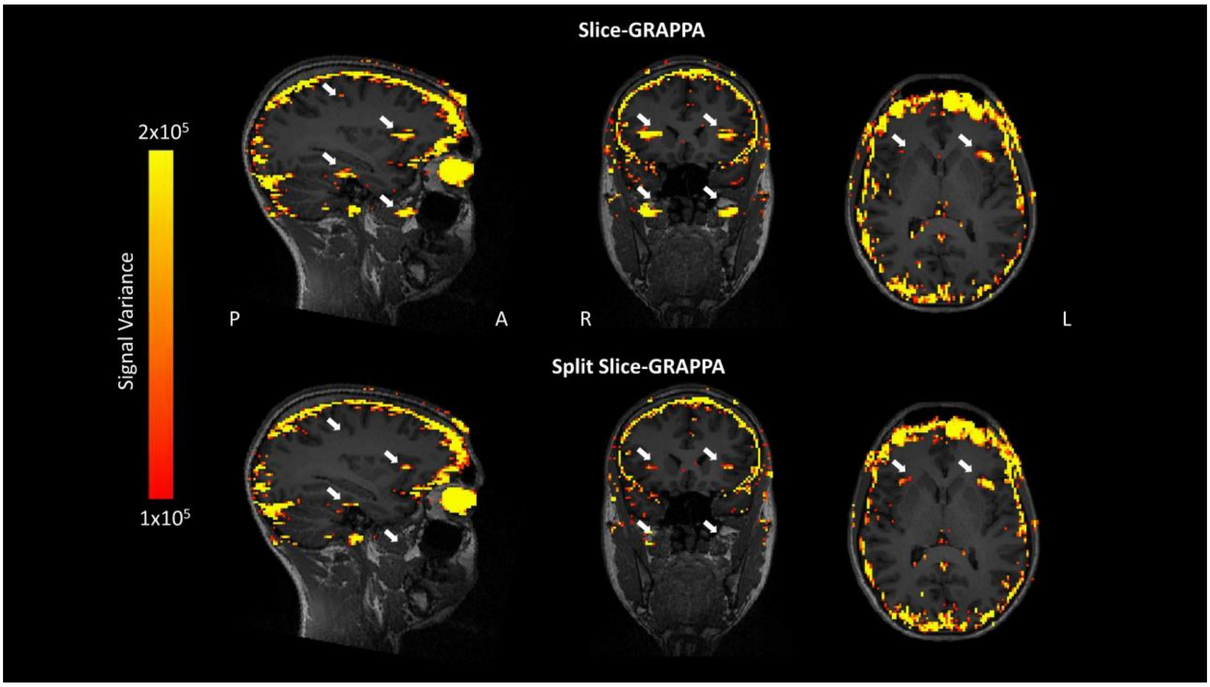


**Fig. S5** Signal variance resulting from slice-GRAPPA (top) and split slice-GRAPPA (bottom) reconstruction techniques using the CMRR multiband acquisition sequence. Data were acquired from two separate acquisitions within an individual participant. Signal variance is in arbitrary units. White arrows indicate expected artefact locations. In-plane CAIPI shift for the CMRR multiband sequence with MB4 and GRAPPA 2 was FOV/4 compared to FOV/3 with Siemens SMS sequence (therefore, artefact locations differ slightly from those presented in previous figures)
